# Macrophages respond rapidly to ototoxic injury of lateral line hair cells but are not required for hair cell regeneration

**DOI:** 10.1101/2020.09.28.314922

**Authors:** Mark E. Warchol, Angela Schrader, Lavinia Sheets

## Abstract

The sensory organs of the inner ear contain resident populations of macrophages, which are recruited to sites of cellular injury. Such macrophages are known to phagocytose the debris of dying cells but the full role of macrophages in otic pathology is not understood. Lateral line neuromasts of zebrafish contain hair cells similar to those in the inner ear, and the optical clarity of larval zebrafish permits direct imaging of cellular interactions. In this study, we used larval zebrafish to characterize the response of macrophages to ototoxic injury of lateral line hair cells. Macrophages migrated into neuromasts within 20 min of exposure to the ototoxic antibiotic neomycin. The number of macrophages in close proximity of injured neuromasts was similar to that observed near uninjured neuromasts, suggesting that this early inflammatory response was mediated by ‘local’ macrophages. Upon entering injured neuromasts, macrophages actively phagocytosed hair cell debris. Such phagocytosis was significantly reduced by inhibiting Src-family kinases. Using chemical-genetic ablation of macrophages prior to ototoxic injury, we also examined whether macrophages were essential for the initiation of hair cell regeneration after neomycin exposure. Results revealed only minor differences in hair cell recovery in macrophage-depleted vs. control fish, suggesting that macrophages are not essential for the regeneration of lateral line hair cells.

## Introduction

Sensory hair cells of the vertebrate inner ear are mechanoreceptors that detect sound vibrations and head movements. Hair cells are also present in the lateral line systems of fish and amphibians, where they detect water motion along the external surface of the animal and convey information to the brain via afferents of the lateral line ganglion (Kindt and Sheets, 2018). Lateral line hair cells are contained in sensory organs known as neuromasts, and are enclosed by an oval-shaped cluster of supporting cells. Because of their optical clarity and well-characterized genetics, the lateral line of larval zebrafish has become a commonly-studied model of hair cell development, pathology and regeneration (Picket and Raible, 2019).

Hair cells of the inner ear can be damaged or lost after noise exposure, ototoxicity or as part of normal aging. Like their counterparts in the ear, lateral line hair cells can also be damaged by exposure to ototoxic drugs or by mechanical trauma (Harris et al., 2003; Hernandez et al., 2006; Uribe et a l., 2018; Holmgren et al., 2020). Many of the signaling pathways that mediate hair cell death have been identified, raising the possibility that some forms of hearing loss may be partially preventable (e.g., Wagner and Shin, 2019). However, the mammalian inner ear cannot regenerate hair cells after injury, and their loss typically results in sensorineural deafness and disequilibrium. In contrast, the ears of nonmammalian vertebrates are able to regenerate hair cells (e.g., Warchol, 2011), and a similar form of hair cell regeneration also occurs in lateral line neuromasts (e.g., Jones and Corwin, 1996; Harris et al., 2003; Denans et al., 2019). At present, the cellular mechanisms that underly hair cell regeneration in nonmammalian vertebrates are poorly understood.

Macrophages are key effector cells of the innate immune system and also contribute to debris clearance and cellular repair after tissue injury. The inner ears of birds and mammals contain resident populations of macrophages, which are recruited to sites of hair cell loss (Warchol, 1997; Bhave et al., 1998; Hirose et al., 2005; Warchol et al., 2012; Kaur et al., 2015). Such macrophages have been shown to phagocytose dying hair cells (e.g., Kaur et al., 2015a; Kaur et al., 2018), but the full role of macrophages in the process of otic pathology is not clear. In addition, the signals that recruit macrophages toward sites of injury in the ear have not been identified (reviewed by Warchol, 2019). The mammalian inner ear is enclosed within the temporal bone of the skull, so it is not possible to directly image the interactions between macrophages and injured hair cells. Notably, however, macrophages are also activated after injury to lateral line neuromasts of fish and salamanders, and their response can be directly imaged in living animals (e.g., Corwin and Jones, 1995; 1996).

The present study used larval zebrafish to characterize the response of macrophages to ototoxic injury of lateral line hair cells. We show that posterior neuromasts typically possess 1-2 macrophages within a 25 μm radius. These macrophages quickly migrate into injured neuromasts during the early phases of ototoxic injury, where they contact and phagocytose the debris of dying hair cells. Although the signals that mediate this response are not fully known, we show that ototoxicity causes the swift externalization of phosphatidylserine on the apical surfaces of hair cells and that macrophage entry into injured neuromasts is reduced after pharmacological inhibition of Src-family kinases. Zebrafish are also capable of regenerating hair cells after ototoxic injury, and we tested whether macrophages were essential for this regenerative response. We found that selective chemical-genetic elimination of macrophages did not affect the extent of ototoxic injury and had only a minor impact on the numbers of regenerated hair cells. Together, these data suggest that macrophages serve as phagocytes after hair cell injury, but do not play a critical role in the initiation of hair cell regeneration.

## Results

### Macrophages reside in close proximity to lateral line neuromasts

Initial studies characterized the association of macrophages with lateral line neuromasts in untreated fish i.e. fish without neuromast injury. Confocal imaging of Tg(*mpeg1:yfp*) zebrafish at 6-7 days-post fertilization (dpf) revealed numerous macrophages distributed throughout the body of each fish. Quantification from low magnification images indicated that the posterior-most 500 μm of larval fish contained ~15 macrophages, many of which were located near neuromasts (Fig, 1A, A’, arrows). Higher magnification images of individual neuromasts showed that most neuromast-associated macrophages possessed a ‘ramified’ morphology, with several processes (pseudopodia) projecting in various directions from the central cell body (Fig. 1B, B’, arrows). Macrophages with a simpler ‘ameboid’ morphology were also observed (Fig. 1B’, B”), but were less common than ramified macrophages. We quantified macrophages located within a 25 μm radius of neuromasts L4 or L5 of the posterior lateral line (nomenclature described in Kindt and Sheets, 2018). Data indicate that, at 6 dpf, neuromasts L4/5 normally possess 1.1±0.7 nearby macrophages (n=15 fish). These observations suggest macrophages may monitor posterior lateral line neuromasts, even in the absence of injury.

**Figure 1.**
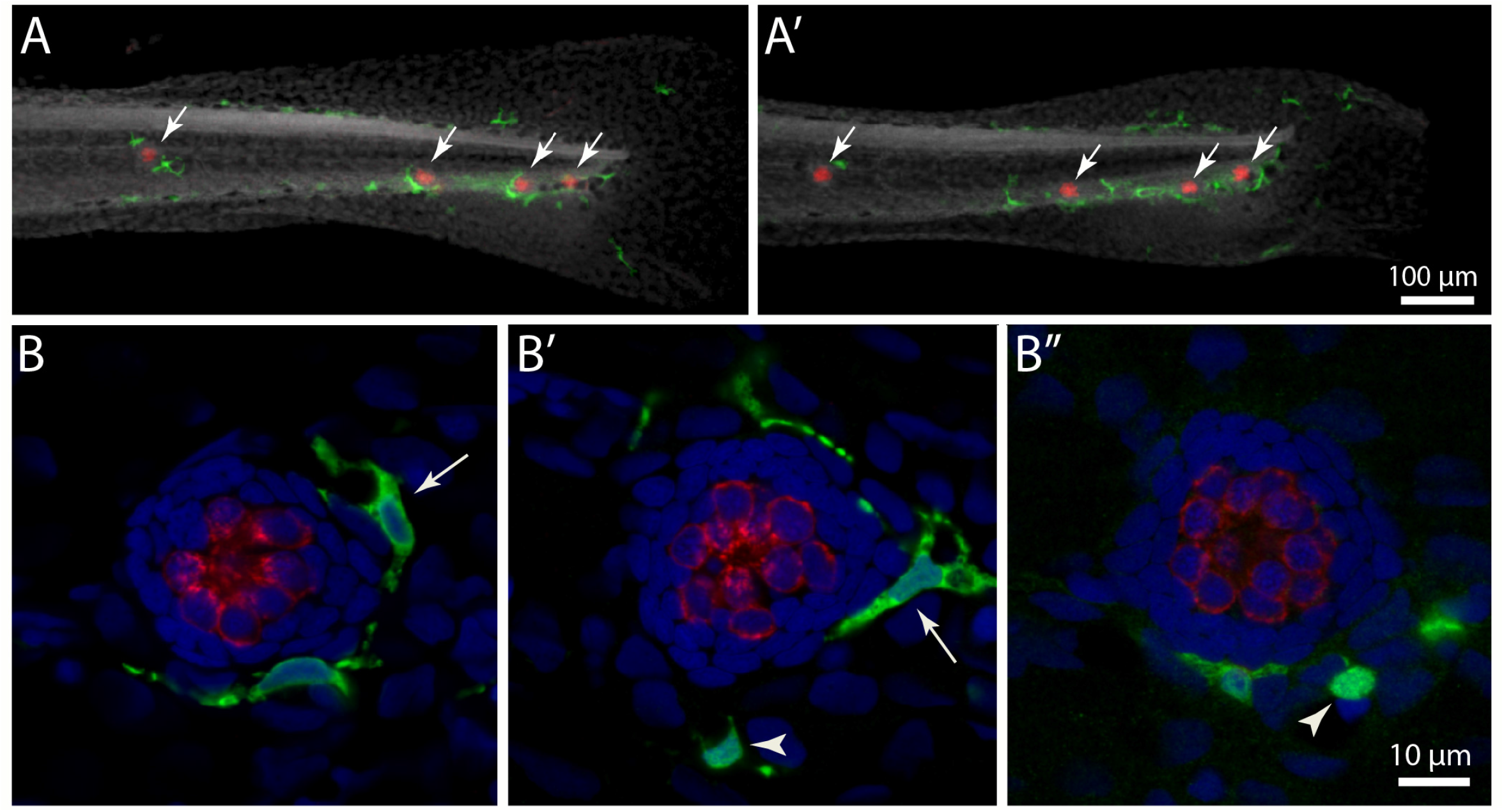
Distribution of macrophages in the posterior lateral line of larval zebrafish. (A, A’):Tail regions of twoTg(mpeg1:yfp) zebrafish, fixed at 6 dpf. YFP in macrophages is shown in green, hair cells are labeled for otoferlin (red), and all nuclei are stained with DAPI (grey). Arrows show the four posterior-most neuromasts in each fish. Note that each neuromast possesses 1-2 nearby macrophages. (B, B’, B”): High magnification views of neuromast L5 from three zebrafish at 6 dpf. Macrophages (arrows) were typically positioned just outside the boundaries of uninjured neuromasts. Most macrophages were elongated and ramified, with 2-3 pseudopodial processes (arrow, B). A smaller number of macrophages with an ameboid morphology were also observed (arrowheads, B’, B”).

### Macrophages contact hair cells after ototoxic injury

In addition to their role in innate immunity, macrophages are also recruited to sites of tissue injury, where they remove the debris of dead cells and secrete bioactive factors that promote repair (e.g., Wynn and Vannella, 2016). We have previously shown that neomycin-induced hair cell death in the posterior-most (‘terminal’) neuromasts of larval zebrafish leads to macrophage entry and phagocytosis (Hirose et al., 2017). A comparable macrophage response was observed in neuromasts L4/5 after exposure to neomycin (Fig. 2). Macrophages typically entered neuromasts within 10-20 min after the initiation of neomycin treatment, and confocal images frequently showed macrophages extending pseudopodial processes that enclosed hair cells (Fig. 2, arrows in 10 min and 20 min images). In addition, many macrophages had internalized otoferlin-labeled material (Fig. 2, arrowhead in 20 min image), which was interpreted as evidence for macrophage phagocytosis of hair cell debris.

**Figure 2.**
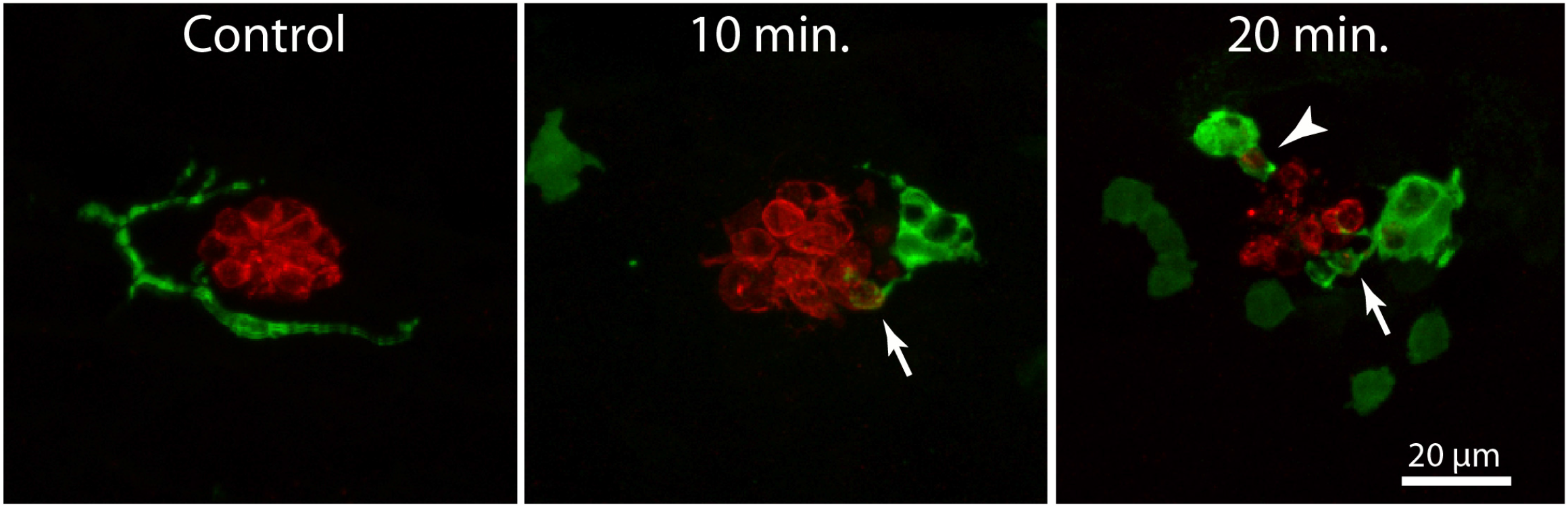
Macrophage response to neomycin ototoxicity. Images are maximum-intensity projections of confocal z-stacks of 15 μm depth. Uninjured neuromasts (‘Control’) usually possess 1-2 adjacent macrophages (green). Macrophages enter the neuromasts shortly after initiating treatment in 100 μM neomycin. Images in center and right show macrophages at 10 and 20 min of exposure to neomycin. Some macrophages (arrows, 10 and 20 min) extend processes that contact hair cells (red, otoferlin), while other macrophages had internalized the debris of dead hair cells (arrowhead, 20 min.)

### Macrophage response occurs shortly after the onset of neomycin exposure

In the mammalian cochlea, increased numbers of macrophages are observed within ~1-2 days of selective hair cell ablation (Kaur et al., 2015). In order to characterize the latency of the macrophage response to the death of lateral line hair cells, we quantified macrophage activity at neuromasts L4/5 at various time points following the initiation of neomycin exposure (5, 10, 20, or 30 minutes) or after 30 minutes of neomycin exposure followed by 1 or 2 hours recovery (Figs. 3 and 4). Confocal images of neuromasts were obtained from immunolabeled specimens that were fixed at all neomycin exposure/survival times (one neuromast/fish, n=13-26 fish/time point, data obtained from three separate trials). Initial loss of hair cells was noted after 5-10 min of neomycin treatment (Fig. 3, top row), and hair cell numbers were significantly reduced after 10 min (Fig. 4A; p=0.019, one-way ANOVA). Pyknotic nuclei were also observed after 5-10 minutes of neomycin treatment (Fig. 3, arrowheads) and their numbers became significantly elevated at 20 min of neomycin exposure (Fig. 4B; p<0.0001, one-way ANOVA). The numbers of pyknotic nuclei peaked at 30 min of neomycin treatment, and then declined, as dead hair cells were cleared from the neuromasts. Few (~1-2) hair cells were present two hours after neomycin exposure (Fig. 3, bottom row, 4A).

**Figure 3.**
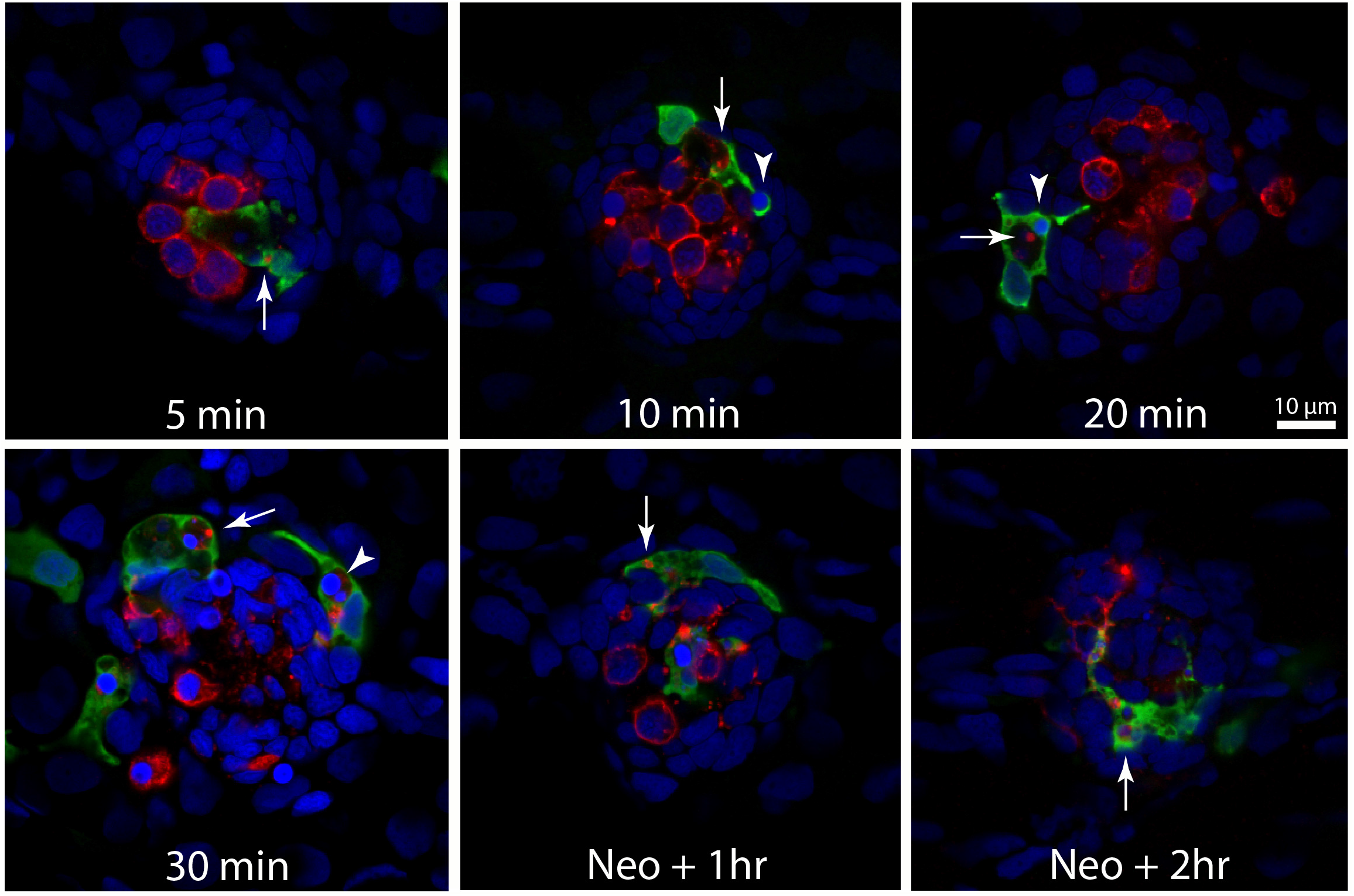
Detailed time course of macrophage response to neomycin ototoxicity. Fish (6 dpf) were incubated in 100 μM neomycin. Some fish were fixed after 5,10,20 or 30 min of neomycin treatment, while other fish were removed from neomycin after 30 min and rinsed and maintained in fresh EM for 1 or 2 hr. Fixed specimens were processed for immunolabeling of YFP-expressing macrophages (green) and hair cells (otoferlin, red), and nuclei were labeled with DAPI (blue). Images are single z-sections taken from 15 μm-depth confocal stacks. Arrows in all images indicate evidence for macrophage phagocytosis of dying hair cells. A few specimens displayed a macrophage response after only 5 min of neomycin exposure. Note that the macrophage in the 5 min image has internalized otoferlin-labeled debris (arrow). At later time points, macrophages had engulfed both otoferlin-labeled debris as well as pyknotic nuclei (arrowheads, 10,20 and 30 min images). Also, beginning at 20 min of neomycin exposure, the number of otoferlin-labeled hair cells is clearly reduced. Neuromasts at 30 min exposure and at 1 and 2 hr recovery typically contained 0-3 surviving hair cells.

**Figure 4.**
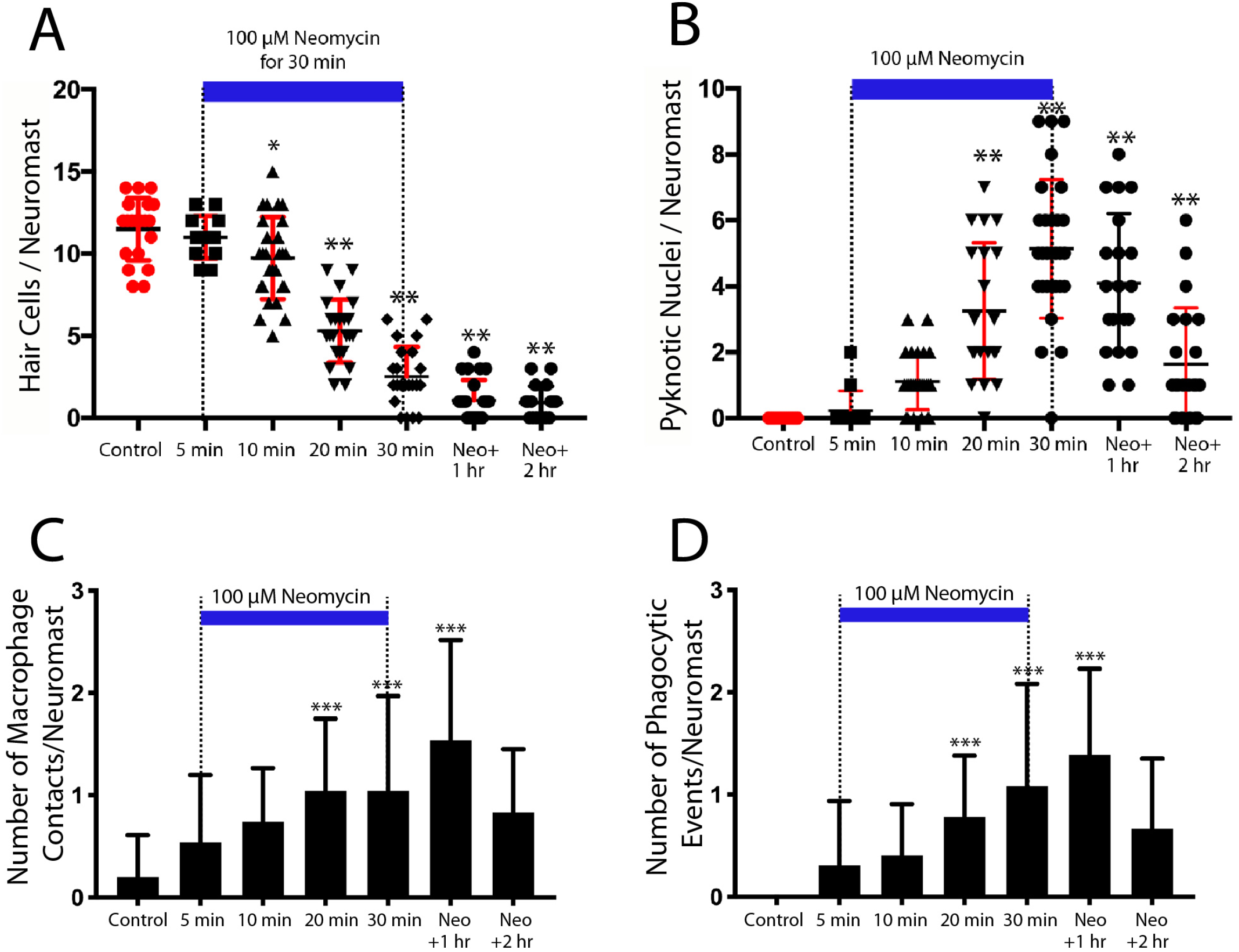
Time course of neomycin ototoxicity and resulting macrophage response. Drug-treated fish were incubated in 100 μM neomycin for various exposure/recovery times, while control fish were incubated in EM for 2 hr. (A) Surviving hair cells as a function of exposure/post-exposure time. Intact hair cells were identified by healthy nuclei that were completely enclosed by otoferlin-labeled hair cell membrane. A significant decrease in hair cell numbers was observed, beginning after 10 min of neomycin exposure (*p=0.019, **p<0.0001, Tukey’s multiple comparisons test). (B) Pyknotic nuclei/neuromast, as a function of neomycin exposure/post-exposure time.The number of pyknotic nuclei/neuromast in neomycin-treated fish was statistically greater than in control neuromasts beginning after 20 min of exposure (**p<0.0001). Although many pyknotic nuclei could not be definitively identified as belonging to hair cells, their number and time course suggests that they represent neomycin-induced death of neuromast hair cells. (C) Contacts between macrophages and hair cells increased after 20 min of neomycin treatment (p=0.0038, one-way ANOVA) and remained elevated until 60 min after treatment. (D) Macrophage phagocytosis of hair cell debris (i.e., internalization of immunolabeled hair cell debris and pyknotic nuclei) was increased, beginning after 20 min of neomycin treatment (p<0.0044, one-way ANOVA).

Macrophages began to enter neuromasts and contact hair cells as early as 5 min into the neomycin treatment (Fig. 3, top left), although such cases were rare. The increase in macrophage contacts with hair cells had significantly increased after 20 min of neomycin treatment and remained enhanced until 60 min after treatment (Fig. 4C, p=0.0038, one-way ANOVA). The numbers of macrophages engaged in phagocytosis (as determined by internalization of either immunolabeled hair cell debris or pyknotic nuclei) also increased after 20 min of neomycin treatment (Fig. 4D, p=0.0044, one-way ANOVA). Despite these changes in macrophage localization and activity, the number of macrophages located within 25 μm of L4/5 at each time point ranged from 1.3-2.0, and did not differ significantly between time points or from controls (p>0.25, one-way ANOVA). This observation suggests that the macrophage response to neomycin injury was mainly carried-out by nearby macrophages, rather than by macrophages recruited from more distant regions of the fish.

### Neomycin treatment leads to rapid externalization of phosphatidylserine

In many tissues, cells undergoing apoptosis are targeted for phagocytic removal via the presence of phosphatidylserine (PtS) on the external membrane surface (Fadok et al., 1992). A prior study has shown external translocation of PtS on the apical surfaces of mouse cochlear hair cells after short exposures to aminoglycoside antibiotics (Goodyear et al., 2006). To determine whether a similar response occurs in zebrafish hair cells, we treated fish in Alexa 555-conjugated annexin V, which binds to and labels externalized PtS (Koopman et al., 1994). Fish were preincubated for 10 min in DAPI (which passes through transduction channels and labels the nuclei of hair cells), and then transferred to embryo medium (EM) containing annexin V. Neomycin was added for a final concentration of 100 μM, and fish were removed, euthanized and fixed, beginning 90 sec after the initiation of neomycin treatment and continuing until 10 min into the treatment. Control fish were treated for 10 min in annexin V but did not receive neomycin. Fluorescent labeling on the apical surfaces of hair cells was observed in nearly all neomycin-treated fish (Fig. 5 arrows), and the numbers of annexin V-labeled hair cells increased during the 10 min exposure (data collected from L4/5; n=17-40 neuromasts/time point). Reconstruction of high magnification images confirmed that the annexin-V label was confined to the apical surfaces of hair cells (Fig. 5, ‘side view’). Quantification of the number of labeled hair cells revealed an increase with longer exposure times (see plot in Fig. 5). In contrast, we observed very few labeled hair cells in fish that were maintained for 10 min in annexin V but not exposed to neomycin (0.7±1.4 HC/neuromast). Notably, annexin V labeling was restricted to the apical (external) surface of hair cells and was not observed along their basal membranes (which are located beneath the epidermis). We also did not observe Annexin V labeling in any cells within the body of any fish, suggesting that Annexin V did not cross the epidermis.

**Figure 5.**
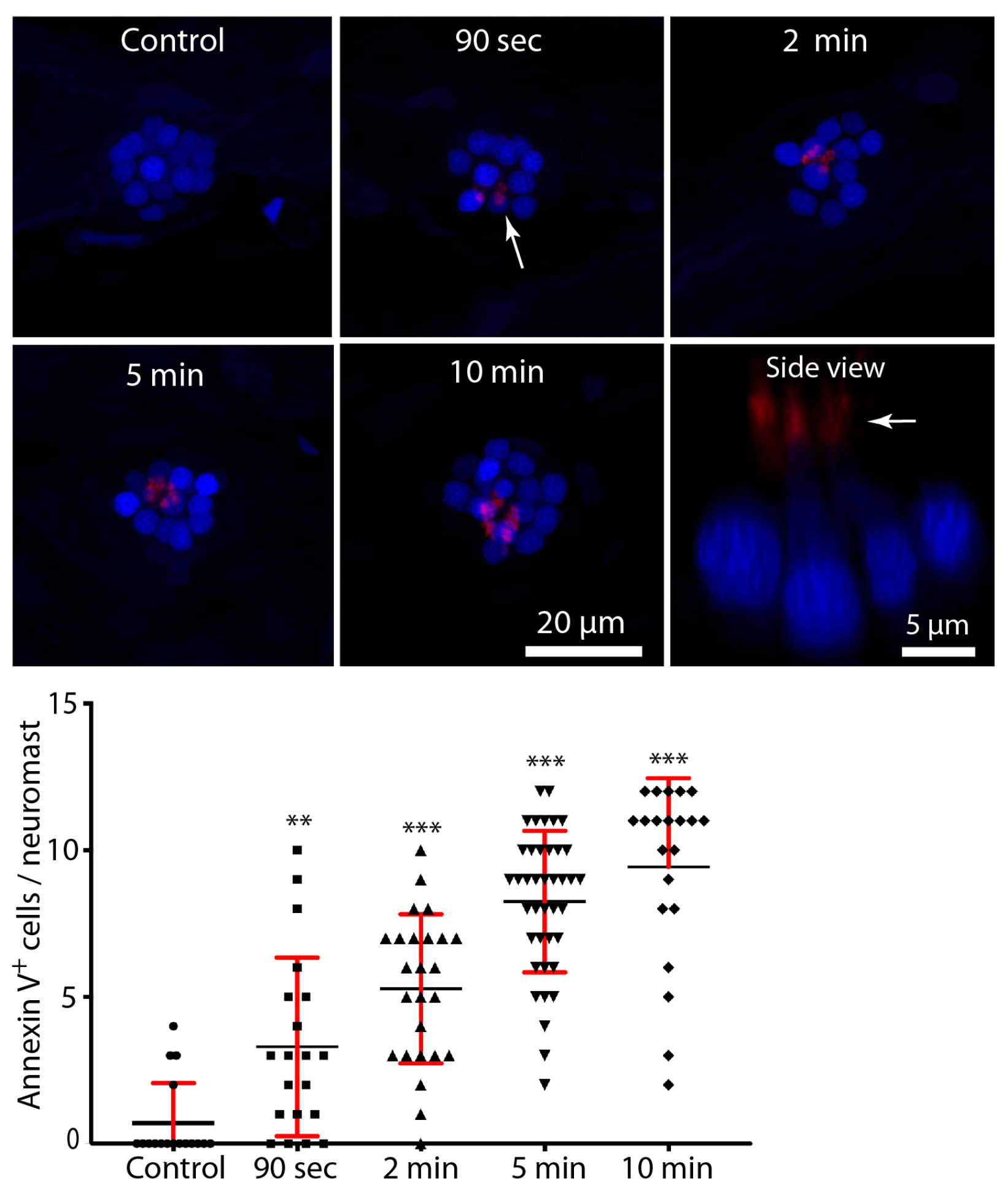
Rapid externalization of phosphatidylserine (PtS) in response to neomycin treatment. Larval zebrafish (6 dpf) were incubated in Alexa 555-conjugated annexin V and neomycin was added to the water, for a final concentration of 100 μM. Fish were euthanized and fixed after 90 sec, 2, 5, or 10 min of exposure to neomycin. Binding of annexin V (which indicates the presence of PtS on the outer membrane surface) was observed as early as 90 sec after the initiation of neomycin treatment (arrow, 90 sec image, p=0.0045, one-way ANOVA), and the number of annexin V-labeled cells increased with longer exposure times (p<0.0001, one-way ANOVA). Annexin V labeling was limited to the apical (external) surface of hair cells, as confirmed by side view reconstruction of confocal stacks (arrow,’side view’). We observed no labeling on the basal hair cell surface, which is located within the epidermal barrier of the fish. We also did not observe annexin V binding on any other cells within the body of the fish, suggesting that annexin V does not become internalized. Plotted data were collected from neuromasts L4/5, for a total of 17-40 neuromasts/time point.

### Inhibition of Src-family kinases reduces macrophage injury response

The signals that trigger the macrophage response to hair cell injury are not known, but activation of Src-family kinases (SFKs) is an evolutionarily-conserved signal known to regulate the activity of phagocytes (Yoo et al., 2012; Freedman et al., 2015; Dwyer et al., 2016). To test whether SFK activation was necessary for macrophage entry into injured neuromasts, we treated fish with PP2, a specific inhibitor of SFKs (Zhu et al., 1999). In initial experiments, fish were treated for 1 hr in either 20 μM PP2 (n=14) or 0.1% DMSO (n=10). Fish were then euthanized and processed for visualization of macrophages and hair cells. No differences were observed in either resident macrophages or hair cell numbers in the PP2-treated vs. control fish (Fig. 6A, data not shown), suggesting that the normal association of macrophages with undamaged neuromasts (e.g., Fig. 1) is not dependent on SFK signaling. We then tested whether SFK signaling affected the macrophage response to hair cell injury by preincubating fish for 1 hr in PP2 or DMSO, and then exposing both groups for 30 min to 100 μM neomycin (n=34 fish/group). The numbers of macrophages in the vicinity of injured neuromasts was not changed by PP2 treatment (Fig. 6D). However, PP2 treatment led to reduced macrophage contact with injured neuromasts following neomycin treatment (Fig 6B, C, E) as well as a reduction in the number of phagocytic events (Fig. 6B, C, F). Together, these observations suggest that SFK signaling contributes to the macrophage response to hair cell injury.

**Figure 6.**
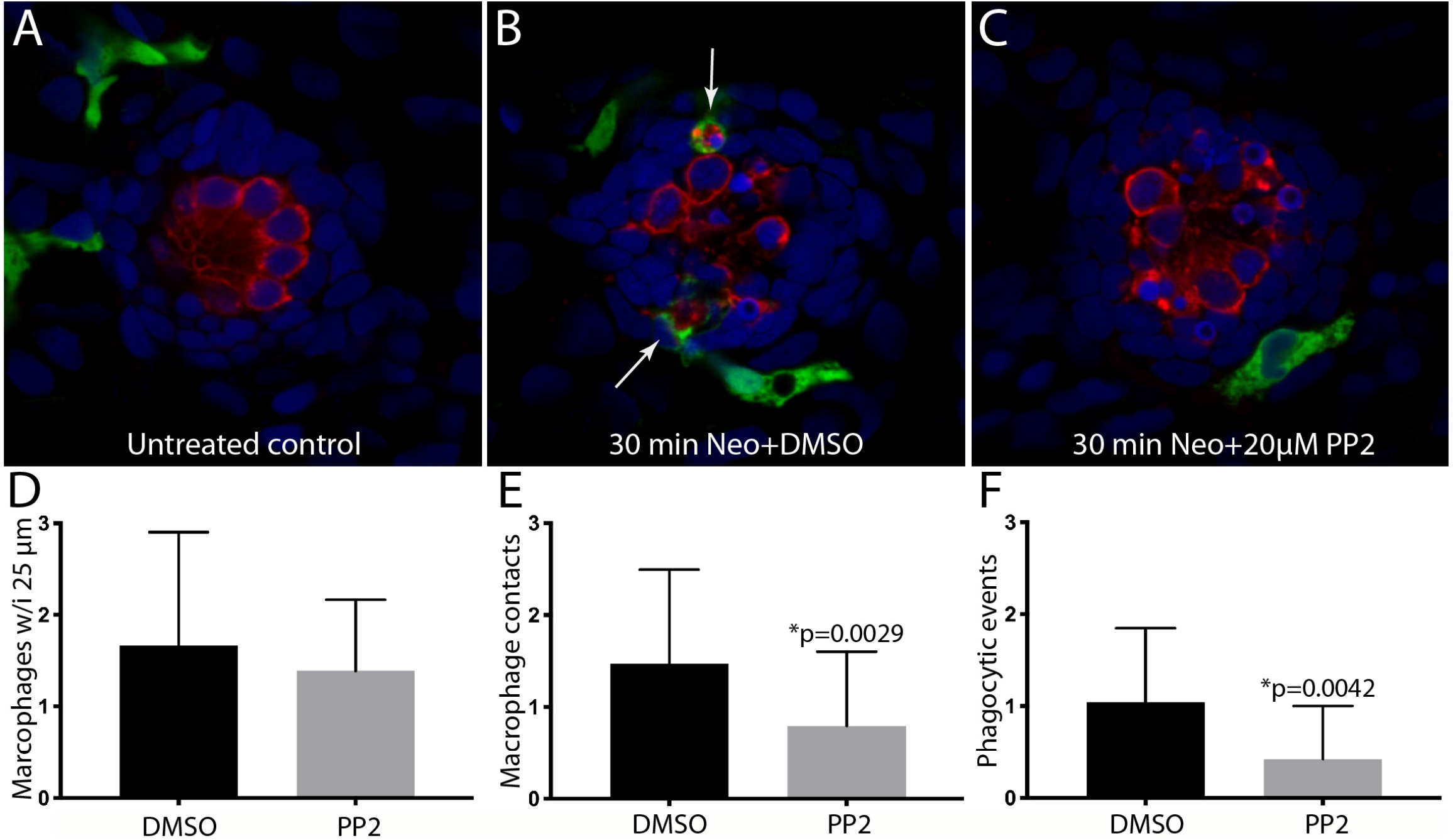
Inhibition of Src-family kinases reduced macrophage entry into injured neuromasts. Larval zebrafish were treated in 20 μM PP2, an inhibitor of several Src kinases. Control fish were treated in parallel with 0.1 % DMSO. After 2 hr pretreatment, neomycin was added to the water of both treatment groups (for a final concentration of 100 μM), and all specimens were euthanized and fixed after 30 min of neomycin exposure. (A-C) Single z-section images taken from 15 μm-depth confocal stacks. Arrows in (B) indicate macrophage phagocytosis of dying hair cells. (D-F) Quantification of macrophage activity. Normal numbers of macrophages were present near neuromasts L4/5 in all fish (D). Fish that were treated with 0.1 % DMSO displayed a normal macrophage response to neomycin injury, but pretreatment with PP2 resulted in fewer macrophage contacts with dying hair cells (E) and reduced numbers of phagocytic events (F). Data were obtained from 34 fish for each treatment group/time point.

### Selective depletion of macrophages has minimal effect on hair cell regeneration

Lateral line neuromasts are able to quickly regenerate hair cells after ototoxic injury, but the biological basis of this regenerative process is not fully understood (Harris et al., 2003; Ma et al., 2008; Romero-Carvajal et al., 2015; Denans et al., 2019). Macrophages have been shown to play an important role in the initiation of regeneration in numerous other tissues and organ systems (reviewed by Keightley et al., 2014; Winn and Vannella, 2016), and it has been suggested that macrophages may be one factor that simulates the production of replacement hair cells in nonmammalian vertebrates (Corwin et al., 1991; Warchol, 1997; Carrillo et al., 2016; Denans et al., 2019). To evaluate the possible role of macrophages in the regeneration of lateral line hair cells, we quantified ototoxic injury and hair cell recovery in a transgenic fish line that permits the selective elimination of macrophages.

*Tg(mpeg1:Gal4FF/ UAS:NTR-mCherry)* double-transgenic fish expresses the gene for nitroreductase under regulation of the macrophage-specific *Mpeg1* promoter (Davison et al., 2007; Ellet et al., 2011). Nitroreductase is a bacterial enzyme that generates a cytotoxin when exposed to the antibiotic metronidazole (MTZ). Incubation of these transgenic fish in MTZ results in selective elimination of macrophages and related cells, without other apparent pathology (Petrie et al., 2014). To verify macrophage depletion, we first treated transgenic fish for 24 hr in 10 mM MTZ (n=10) or in 0.1% DMSO (controls, n=15), and then quantified fluorescently-labeled macrophages in posterior-most 500 μm of all fish (Fig. 7). Control fish contained 16.3±4.9 macrophages in this region (Fig 7A, C), but the MTZ-treatment dramatically reduced the number of macrophages, to 0.8±1.1. When MTZ-treated fish were allowed to recover for 48 hr (n=14), the tail contained 3.2±2.2 macrophages (p<0.0001, two-tailed T-test), indicating that significant macrophage depletion persists for (at least) two days. In order to verify that expression of the *Ntr* transgene did not affect the normal development of the lateral line, we also examined hair cell numbers in *Tg(mpeg1:Gal4FF/ UAS:NTR-mCherry)* and sibling fish that lack the transgenes. Neuromasts L4/5 of *Ntr*-expressing fish contained 10.1±1.4 hair cells vs. 10.6±1.2 hair cells in fish that lacked the *Ntr* transgene (n=10/15, p=0.35).

**Figure 7.**
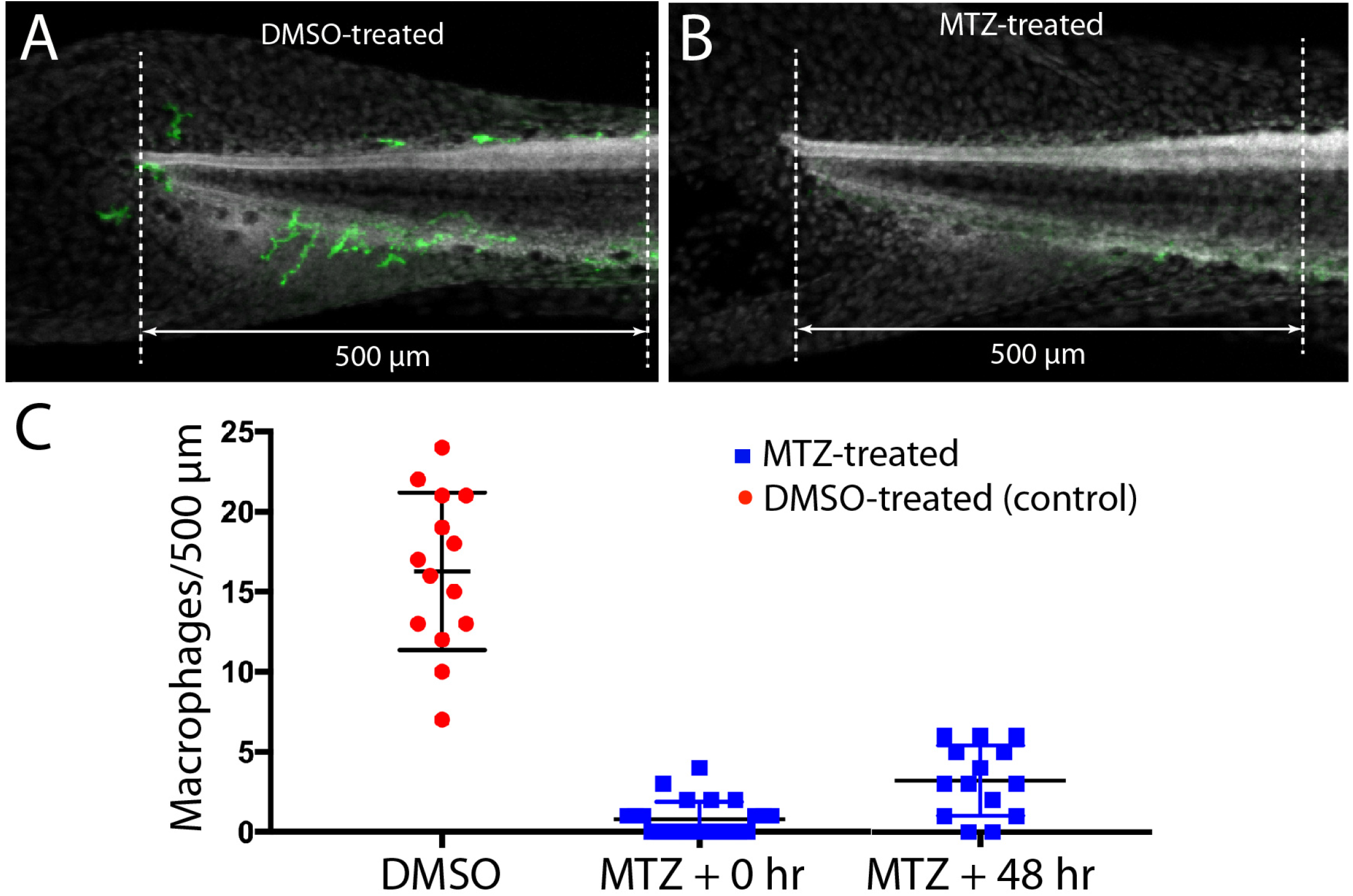
Selective depletion of macrophages in Tg(mpeg1:Gal4FF/ UAS:NTR-mCherry) fish. Experiments utilized a transgenic fish line that expresses the bacterial enzyme NTR under regulation of the Mpeg1 promoter (i.e., selectively in macrophages and microglia). Treatment of these fish with 0.1% DMSO did not affect the numbers of macrophages distributed within the posterior-most 500 μm of the spinal column (A, C). However, treatment for 24 hr in 10 mM MTZ resulted in nearly complete elimination of macrophages from this same region (B, C). Macrophages numbers remained low after 48 hr recovery from MTZ treatment (C). Data obtained from 10-15 fish/treatment group.

To determine whether the depletion of macrophages affected hair cell regeneration after ototoxic injury, *Tg(mpeg1:Gal4FF/ UAS:NTR-mCherry)* fish were treated for 24 hr with either 10 mM MTZ or 0.1 % DMSO. All fish were then rinsed and exposed for 30 min to 100 μM neomycin, at which point they were thoroughly rinsed and returned to drug-free EM. Hair cells were quantified from neuromasts L4/L5 and from the two most-posterior (‘terminal’) neuromasts at 2 hr and 48 hr after neomycin treatment. At 2 hr post-neomycin, the number of surviving hair cells was nearly-identical in both MTZ-treated and control fish (Fig 8C; 1.1±1.2 vs. 0.9±1.0; n=30/40; p=0.32), suggesting that macrophage depletion did not affect ototoxicity. Hair cell regeneration was evident in both MTZ-treated and control fish 48 hr after neomycin treatment. In neuromasts L4/5, hair cell number in MTZ-treated fish was 6.7±2.2 vs. 7.7±2.0 in DMSO-treated fish (n=19/group, p=0.19). In the two terminal neuromasts, the numbers of regenerated hair cells in MTZ-treated fish was 5.6±1.5 vs. 6.9±1.8 in DMSO-treated fish (n=44/38; p=0.0012). Together, these observations suggest that depletion of macrophages had only a small effect on hair cell regeneration after neomycin ototoxicity.

**Figure 8.**
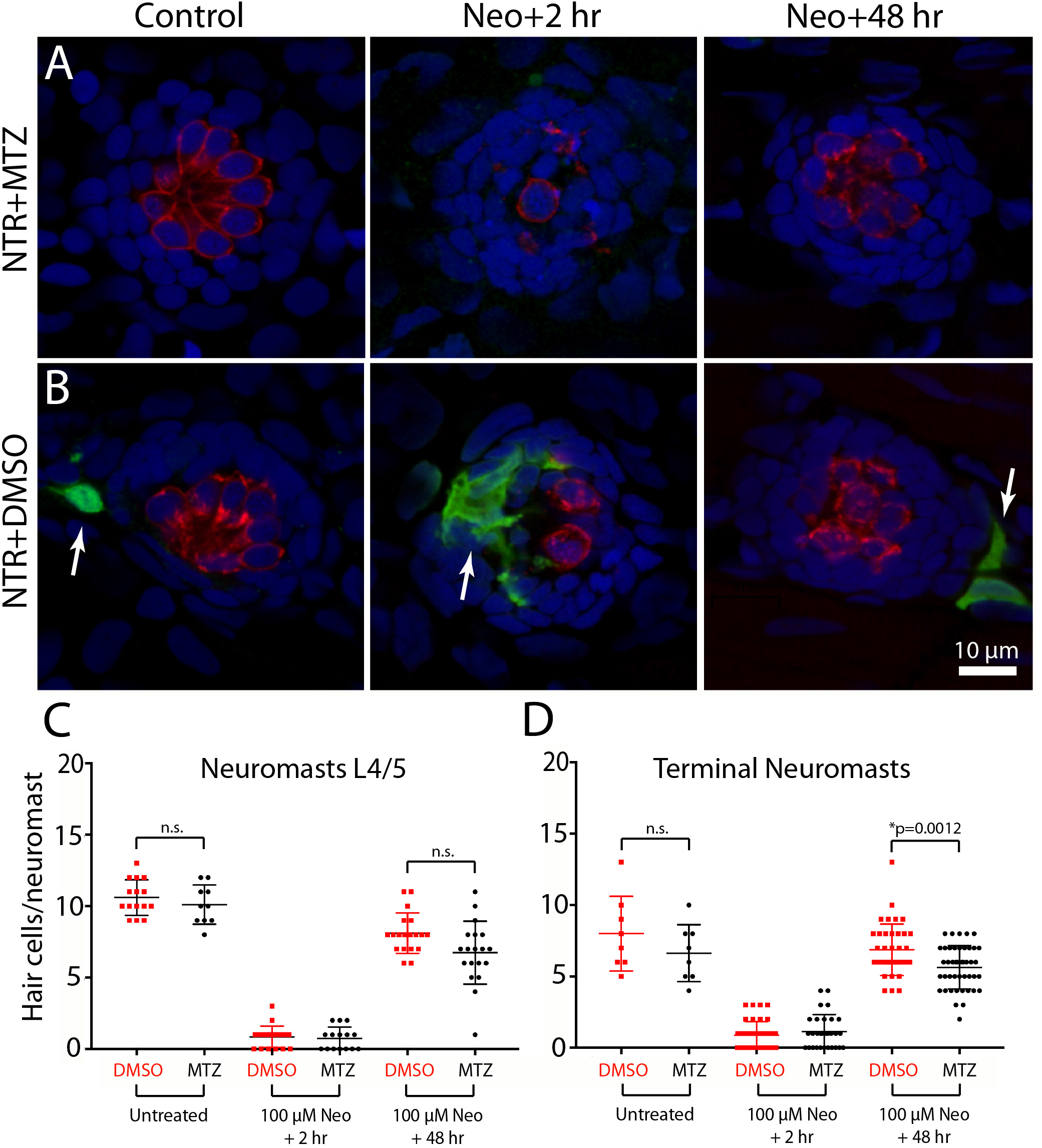
Depletion of macrophages had minimal impact on hair cell regeneration. Tg(mpeg1:Gal4FF/ UAS:NTR-mCherry) fish were treated for 24 hr in either 10 mM MTZ or 0.1% DMSO. All fish were then rinsed and incubated for 30 min in 100 μM neomycin. Fish were rinsed again and maintained for an additional 2 hr or 48 hr. Hair cells and macrophages were immunolabeled and hair cell numbers were quantified from neuromasts L4/5 and the two terminal neuromasts. Images from MTZ-treated fish (A) show absence of macrophages, but otherwise normal levels of neomycin injury and subsequent regeneration. Ototoxicity and regeneration were also unaffected in DMSO-treated fish (B), but macrophage numbers and behavior were normal (arrows). Quantitative data (C, D) show that hair cell injury and regeneration were nearly-identical in normal and macrophage-depleted fish.The number of regenerated hair cells in the two terminal neuromasts of MTZ-treated was slightly reduced, when compared to DMSO-treated controls (p=0.0012)

## Discussion

Macrophages are a type of leukocyte that recognize, engulf and neutralize invading pathogens and are also actively involved in tissue homeostasis and injury response (e.g., Gordon and Pluddemann, 2018). Macrophages are recruited to tissue lesion sites, where they remove the debris of dead cells and release extracellular matrix constituents and growth factors that promote repair and regeneration. The inner ears of birds and mammals contain resident macrophages which are activated by hair cell injury (Warchol, 1997; Bhave et al., 1998; Hirose et al., 2005; Warchol et al., 2012). Prior studies have shown that otic macrophages remove cellular debris after noise trauma or ototoxicity (Kaur et al., 2015b, Kaur et al., 2018). It should be noted, however, that macrophages are not the only resident phagocyte in the inner ear. The supporting cells of the sensory epithelium can also act as ‘amateur phagocytes,’ by forming multi-cellular complexes that engulf dying cells (Bird et al., 2010; Anttonen et al., 2014; Monzack et al., 2015; Bucks et al., 2017; reviewed by Hirose et al., 2017).

Our data suggest that the interactions between macrophages and lateral line hair cells are similar to those that occur in the inner ear. As noted, the uninjured cochlea and vestibular organs possess resident populations of macrophages (Warchol, 1997; Hirose et al., 2005; Kaur et al., 2015b), and our data indicate that 1-2 macrophages are typically present within a 25 μm radius of uninjured neuromasts (Fig. 1; also see Herbomel et al., 1999). The signals responsible for the association between macrophages and hair cell epithelia are not known, but our data are consistent with the notion that a chemoattractant may be released from uninjured neuromasts, so as to maintain a nearby ‘resident’ population. Our data further show that the numbers of macrophages present at injured neuromasts is comparable to the numbers observed at undamaged neuromasts. This result, which is consistent with previous data (Hirose et al., 2017), suggests that the initial macrophage response to hair cell injury is mediated by ‘local’ macrophages.

Studies of the mammalian ear indicate that hair cell death alone – without any accompanying damage to other cells or tissues – is sufficient to recruit macrophages into the sensory regions of the cochlea and utricle (Kaur et al., 2015a, 2015b). The signals that evoke this response have not been identified. Our data show that neomycin-induced hair cell death causes macrophages to enter lateral line neuromasts, and the rapid nature of this response suggests that hair cells (or perhaps supporting cells) release a chemotactic signal early in the ototoxic process. Numerous signals have been shown to recruit macrophages to injury sites in other tissues. These include the pannexin-mediated release of ATP (Adamson and Leitinger, 2014), the generation and release of reactive oxygen species (ROS) such as H_2_O_2_ (Tauzin et al., 2014), and the release of certain chemokines (Gillitzer and Goebeler, 2001). It is further notable that both ATP and ROS are known to be produced during ototoxic injury and hair cell death (e.g., Gale et al., 2004; Lahne and Gale, 2010; Esterberg et al., 2016), making them ideal candidates for future studies of hair cell-macrophage interactions. Moreover, the rapid macrophage response reported here suggests that zebrafish lateral line may be an advantageous model system in which to identify the signals that attract macrophages to injured hair cells.

The signals that target dying hair cells for phagocytosis are also not known. One common ‘eat me’ signal displayed by dying cells is the presence of phosphatidylserine (PtS) on the external membrane surface. Healthy cells possess asymmetrical localization of phospholipids on the inner and outer membrane leaflets, with PtS present on the inner leaflet and phosphatidylcholine exposed on the outer leaflet (Nagata, 2018). This distribution pattern is maintained by ATP-dependent transporters (‘flippases’). As noted, dying cells are characterized by the presence of PtS on the outer membrane leaflet. The internal-to-external translocation of PtS is mediated via two distinct mechanisms: (1) Ca^2+^-dependent activation of the TMEM16 transporter and (2) caspase-dependent activation of the XKR8 ‘scramblase’. Externalized PtS is a highly-conserved signal that is recognized by phagocytes, and leads to the engulfment and removal of apoptotic cells (reviewed by Kloditz et al., 2017). The present study used Alexa 555-conjugated Annexin V to examine PtS externalization in hair cells after neomycin ototoxicity. We found that very few hair cells in normal (control) fish possessed external PtS. However, enhanced levels of externalized PtS were observed on the apical surfaces of lateral line hair cells after only 90 sec of neomycin treatment, and the number of such PtS-expressing cells increased with longer exposure times. These observations are very similar to those reported in studies of the mammalian cochlea (Goodyear et al., 2008), and suggest that neomycin induces very rapid changes in hair cell homeostasis. The timing is also consistent with the notion that externalized PtS may target injured hair cells for phagocytic removal. Notably, macrophages are likely to detect PtS on the basal (internal) surfaces of hair cells, while we observed annexin V labeling only on the apical (external) surface. Given that larval fish are likely to contain numerous cells undergoing developmental apoptosis (e.g., Shklover et al., 2015), we speculate that the lack of internal annexin V labeling is a consequence of the inability of annexin V to cross the epidermal barrier. We cannot, however, rule out the possibility that externalized PtS is not present on the basal surfaces of injured hair cells.

Our data further indicate that activation of Src family kinases is involved in the entry of macrophages into injured neuromasts and the subsequent phagocytosis of hair cell debris. The Src family consists of nine nonreceptor tyrosine kinases that serve diverse roles in cell signaling. Macrophages express several SFK’s (e.g., Fgr, Fyn, Hck, Lyn), which are activated downstream of toll-like receptors and regulate macrophage response to pathogens (reviewed by Byeon et al., 2012). In addition, Lyn can be directly activated by diffusible H_2_O_2_, which can serve as a leukocyte chemoattractant (Yoo et al., 2011; 2012). A prior study has shown that ROS are generated by zebrafish hair cells within 5-10 min of exposure to neomycin (Esterberg et al., 2016), raising the possibility that such hair cells may release H_2_O_2_, and that this might recruit macrophages into the neuromast. Generation of ROS also occurs in mammalian hair cells after noise trauma or ototoxic injury (reviewed by Fetoni et al., 2019) and it conceivable that extracellular release of ROS may also promote macrophage migration towards the injured organ of Corti. In future studies, the possible involvement of Lyn activation and SFK signaling in hair cell-macrophage interactions in the mammalian ear can be tested using genetic knockout animals (e.g., Chan et al., 1997).

Finally, our experiments tested the possible role of macrophages in the process of hair cell regeneration. Nonmammalian vertebrates can regenerate hair cells after injury, resulting in the restoration of sensory function (e.g., Warchol, 2011). Such regeneration is mediated by nonsensory supporting cells, which surround hair cells in their native epithelia and are involved in homeostasis and regulation of the ionic environment. Hair cell death in nonmammals triggers supporting cells to either divide or undergo phenotypic conversion, leading to the production of replacement hair cells (reviewed by Warchol, 2011; Denans et al., 2019). The biological signals responsible for hair cell regeneration are largely unidentified. Given that macrophages have been shown to promote regeneration in many injured tissues (e.g., Theret et al., 2019), it has been proposed that recruited macrophages may also help initiate hair cell regeneration (e.g., Corwin et al., 1991; Warchol, 1997; 1999). The rapid time course of macrophage recruitment is consistent with this notion, since macrophages migrate to sites of hair cell injury prior to the onset of regenerative proliferation or phenotypic conversion (Warchol, 1997; Hirose et al., 2017). However, a more recent study of regeneration in the avian cochlea found that the selective macrophage depletion had no impact on either supporting cell proliferation or hair cell replacement (Warchol et al., 2012). The data presented here further indicate that macrophages are not essential for hair cell regeneration. Macrophages quickly entered injured neuromasts and phagocytosed dying cells, but chemical-genetic depletion of macrophages had very little impact on the level of hair cell regeneration after neomycin injury. In addition, macrophage depletion did not affect the number of surviving hair cells or the extent of ototoxic injury. It should be noted that Warchol et al. (2012) found that depletion of macrophages caused a reduction in proliferation of some mesenchymal cells that reside outside of the sensory epithelium, so it is still possible that macrophages may be involved in the maintenance and repair of nonsensory structures within the inner ear.

A prior study has also examined the contribution of macrophages to hair cell regeneration in larval zebrafish (Carrillo et al., 2016). Macrophage depletion was induced by local injections of liposomally-encapsulated clodronate, which reduced macrophage numbers in the vicinity of identified neuromasts. These interventions appeared to cause a slight reduction in the number of regenerated hair cells/neuromast at 24 hr recovery (~4 HCs in clodronate-treated vs. ~5 HCs in controls), but no difference in regeneration at 48 hr recovery. The data reported by Carrillo et al., (2016) are generally consistent with those described reported here, i.e., we found that depleting macrophages caused a slight reduction in regeneration in the terminal neuromasts. However, the findings of both studies demonstrate that the presence of normal numbers of macrophages is not necessary for lateral line regeneration.

In summary, our results show that macrophages normally reside near lateral line neuromasts and are recruited into neuromasts in the early stages of ototoxic injury. Such macrophages quickly engulf the debris of dying hair cells. Our data further suggest that activation of Src-family kinases is required for normal macrophage recruitment and that dying hair cells possess externalized PtS, a signal that targets other types of apoptotic cells for phagocytic removal. However, our data indicate that selective elimination of macrophages had a very minimal effect on hair cell recovery after ototoxic injury, suggesting that macrophages do not serve an essential role in the process of hair cell regeneration. The molecules that mediate signaling between sensory hair cells and macrophages are currently not known, but our findings suggest that the zebrafish lateral line may be an advantageous model system in which to study hair cell-immune interactions.

## Methods

### Ethics Statement

This study was performed with the approval of the Institutional Animal Care and Use Committee of Washington University School of Medicine in St. Louis and in accordance with NIH guidelines for use of zebrafish.

### Zebrafish

Most experiments were performed using Tg(*mpeg1:yfp*) fish, which selectively express YFP under regulation of the *Mpeg1* promoter (i.e., in macrophages and microglia – Ellett et al., 2011; Svahn et al., 2013; Roca & Ramakrishnan, 2013). Studies of hair cell regeneration used *Tg(mpeg1:Gal4FF/ UAS:NTR-mCherry)* double transgenic fish, which express the Gal4 transcriptional activator driven by the macrophage-specific *Mpeg1* promoter and the gene for for the bacterial enzyme nitroreductase fused to mCherry under regulation of the Gal4-specific UAS enhancer sequence. Adult zebrafish were maintained at 27-29°C and housed in the Washington University Zebrafish Facility. Fertile eggs and larvae were maintained in embryo medium (EM: 15mM NaCl, 0.5mM KCl, 1mM CaCl_2_, 1mM MgSO_4_, 0.15mM KH_2_PO_4_, 0.042 mM Na_2_HPO_4_, 0.714mM NaHCO_3_ (Westerfield, 1994) and, beginning at 5 dpf, were fed rotifers daily. At the end of experiments, fish were euthanized by quick chilling to 4° C.

### Ototoxic ablation of neuromast hair cells with neomycin

Lateral line hair cells were lesioned by incubating fish in the ototoxic antibiotic neomycin (e.g., Harris et al., 2003). Larval fish were placed in 25 mm ‘baskets’ (Corning Cell Strainer, ~20-30 fish/basket) and transferred into 30 ml EM that contained 100 μM neomycin (Sigma). Depending on the specific experiment, fish were treated in neomycin for 90 sec-30 min, and were then either euthanized and fixed, or rinsed 3x by immersion in 30 ml EM and maintained for an additional 1-48 hr.

### Treatment with SFK inhibitor

To examine the influence of inhibiting Src-family kinases on macrophage response to ototoxic injury, fish were treated in PP2, an inhibitor of SFKs (Caymen Chemical, 20 μM). A 20 mM stock solution was prepared in DMSO and diluted 1:1,000 in EM. Control specimens were maintained in parallel in 0.1% DMSO.

### Selective depletion of macrophages

The influence of macrophages on hair cell regeneration was examined using *Tg(mpeg1:Gal4FF/ UAS:NTR-mCherry)*. Macrophages were eliminated via incubation for 24 hr in 10 mM MTZ (Sigma, with 0.1% DMSO). Controls in these studies were fish of the same genotype, but incubated 24 hr in 0.1% DMSO alone.

### Immunohistochemical labeling

Following overnight fixation at 4°C, fish were thoroughly rinsed in PBS, and nonspecific antibody binding was blocked by treatment for 2 hr in 5% normal horse serum (NHS) in phosphate-buffered saline (PBS) with 1% Triton X-100. This was followed by incubation in primary antibodies, which were diluted in PBS, with 2% NHS and 1% Triton X-100. All specimens were treated in antibody solutions overnight, at room temperature and with mild agitation. The next day, specimens were rinsed 3x in PBS and incubated for 2 hr in secondary antibodies (anti-mouse IgG and anti-rabbit IgG, both raised in donkey) that were conjugated to Alexa-488 and Alexa-555, respectively (1:500, Invitrogen). The secondary solution also contained the nuclear dye DAPI. Following thorough rinsing in PBS, fish were mounted in glycerol:PBS (9:1) on microscope slides and coverslipped.

### Primary antibodies

Hair cells were labeled with HCS-1, which is specific for otoferlin (Goodyear et al., 2010). HCS-1 was developed by Jeffrey Corwin (University of Virginia) and obtained from the Developmental Studies Hybridoma Bank, created by the NICHD of the NIH and maintained at Department of Biology of the University of Iowa. HCS-1 was obtained as a purified concentrate and used at 1:500 dilution. The YFP or mCherry signals in macrophages were amplified by labeling with either anti-GFP or anti-mCherry (1:500; ThermoFisher).

### Confocal imaging

Images of fixed samples were acquired using an LSM 700 laser scanning confocal microscope (Carl Zeiss). Confocal stacks of 15 μm depth were collected with a z-step of 1 μm. Regardless of the particular fluorophore expressed (either YFP or mCherry), macrophages in all presented images were pseudo-colored green.

### Confocal image processing and analysis

Confocal image stacks were reconstructed and analyzed using Volocity software (Quorum Technologies). Intact hair cells were identified by the presence of a normal-looking DAPI-stained nucleus that was surrounded by an uninterrupted region of otoferlin immunoreactivity. Pyknotic nuclei were identified as small, dense and intense puncta of DAPI labeling. Macrophage activity was quantified by scrolling through the 15 μm-depth image stacks (in the z-dimension) and counting (1) the number of macrophages a within 25 μm radius of a neuromast (using a circle inscribed on the particular neuromast), (2) the number of macrophages directly contacting otoferlin-labeled hair cells, and (3) the number of macrophages that had internalized otoferlin-labeled material (hair cell debris). For each metric, the recorded number reflected the activity of a single macrophage, i.e., a macrophage that made contacts with multiple hair cells and/or had internalized debris from several hair cells was still classified as a single ‘event.’ Subsequent image processing for display as figures was performed using Photoshop and Illustrator software (Adobe).

### Statistical analysis

Plotting and statistical analyses of data were performed using Prism 8 (Graphpad Software Inc). Statistical significance between two (parallel) data sets was determined via unpaired Student’s *t* test or Mann–Whitney U test, as appropriate. Statistical comparison of multiple parallel data sets was determined by one-way ANOVA or Kruskal-Wallis tests, with and appropriate post-hoc tests. All plots show mean±standard deviation.

## Notes

Supported by NIH grant R01DC006283 (M.E.W.) and the WUSTL Department of Otolaryngology (L.S.). Transgenic fish lines were provided by Drs. David Raible and Lalita Ramakrishnan.

### Competing Interest Statement

The authors have declared no competing interest.

